# Using phylogenetic summary statistics for epidemiological inference

**DOI:** 10.1101/2024.08.07.607080

**Authors:** Rafael C. Núñez, Gregory R. Hart, Michael Famulare, Christopher Lorton, Joshua T. Herbeck

## Abstract

Since the coining of the term phylodynamics, the use of phylogenies to understand infectious disease dynamics has steadily increased. As methods for phylodynamics and genomic epidemiology have proliferated and grown more computationally expensive, the epidemiological information they extract has also evolved to better complement what can be learned through traditional epidemiological data. However, for genomic epidemiology to continue to grow, and for the accumulating number of pathogen genetic sequences to fulfill their potential widespread utility, the extraction of epidemiological information from phylogenies needs to be simpler and more efficient. Summary statistics provide a straightforward way of extracting information from a phylogenetic tree, but the relationship between these statistics and epidemiological quantities needs to be better understood. In this work we address this need via simulation. Using two different benchmark scenarios, we evaluate 74 tree summary statistics and their relationship to epidemiological quantities. In addition to evaluating the epidemiological information that can be inferred from each summary statistic, we also assess the computational cost of each statistic. This helps us optimize the selection of summary statistics for specific applications. Our study offers guidelines on essential considerations for designing or choosing summary statistics. The evaluated set of summary statistics, along with additional helpful functions for phylogenetic analysis, is accessible through an open-source Python library. Our research not only illuminates the main characteristics of many tree summary statistics but also provides valuable computational tools for real-world epidemiological analyses. These contributions aim to enhance our understanding of disease spread dynamics and advance the broader utilization of genomic epidemiology in public health efforts.

**Author Summary:** Our study focuses on the use of phylogenetic analysis to get valuable epidemiological insights. We conducted a simulation study to evaluate 74 phylogenetic summary statistics and their relationship to epidemiological quantities, shedding light on the potential of each of these statistics to quantify different characteristics of disease spread dynamics. Additionally, we assessed the computational cost of each statistic. This gives us additional information when selecting a statistic for a particular application. Our research is available through an open-source Python library. This work helps us enhance our understanding of phylogenetic tree structures and contributes to the broader application of genomic epidemiology in public health initiatives.

## 1. Introduction

Phylogenetic analysis plays an increasingly important role in epidemiology and public health, as it can facilitate deep understanding of the spread and evolution of transmissible diseases. Its significance is demonstrated by the pivotal role that phylodynamics and genomic epidemiology has played in unraveling the complex dynamics of various infectious diseases such as HIV (1,2), influenza (3,4), Ebola (5), Mpox (6), and SARS-CoV-2 (7), among many others. The prevailing popularity of phylogenetics can be attributed to multiple factors, including advancements in genome sequencing technology, reduced sequencing costs, faster sequencing times, and the notable expansion of genomic databases.

Despite the increased use of phylogenetic analysis in epidemiology, the development of analytical methods has outpaced the accessibility of such methods. The potential of genomic data is unrealized, due to the scarcity of expertise in handling and analyzing such data, as well as the absence of streamlined and easily usable workflows. These workflows often manifest as subjective processes, rely on the characterization of phylogenetic trees via summary statistics that may be challenging to design, require a linked epidemiological model with accompanying assumptions, or whose application is not entirely clear. For example, researchers might struggle with interpreting the branching patterns of a phylogenetic tree to accurately infer the evolutionary relationships among strains.

The myriad available summary statistics further compound the issue, posing questions such as: Which statistic is the most suitable for my specific application? Do I need to create a new statistic, and if so, what considerations should guide its design and testing? In this paper, we address these questions and, more generally, the need for easier, faster, and increased adoption of phylogenetic analyses, by conducting a thorough evaluation of the performance of numerous summary statistics. Our approach involved a literature search to compile a comprehensive list of summary statistics, followed by an analysis of their performance in two generic epidemic scenarios. Our software to estimate these summary statistics, `phylomodels`, is now available as an open-source Python library, aiming to provide a valuable tool for phylogenetic analysis and epidemiological research.

### 1.1 Related Research

Phylogenetic summary statistics have been studied extensively. These statistics are often introduced with specific objectives in mind, such as quantifying tree imbalances or as an adaptation from other fields like graph or spectral theories. Here, we investigated 74 summary statistics; definitions and references are presented in Table 1. Each of these statistics has a previously demonstrated value for at least one scientific application. Our study delves deeper by examining the performance of these statistics in two benchmark applications and identifying their core strengths (see Table 1). Concurrent to this work, Jenzen & Ettiene performed a similar study of 54 summary statistics and introduced the `Treestats` R package (8). Our study covers most of the same summary statistics as they do while including additional statistics. Their study also leans more toward the taxonomic roots of phylodynamics and focus on diversification and taxonomic groups, while we focus more on transmission dynamics.

**Table 1.**
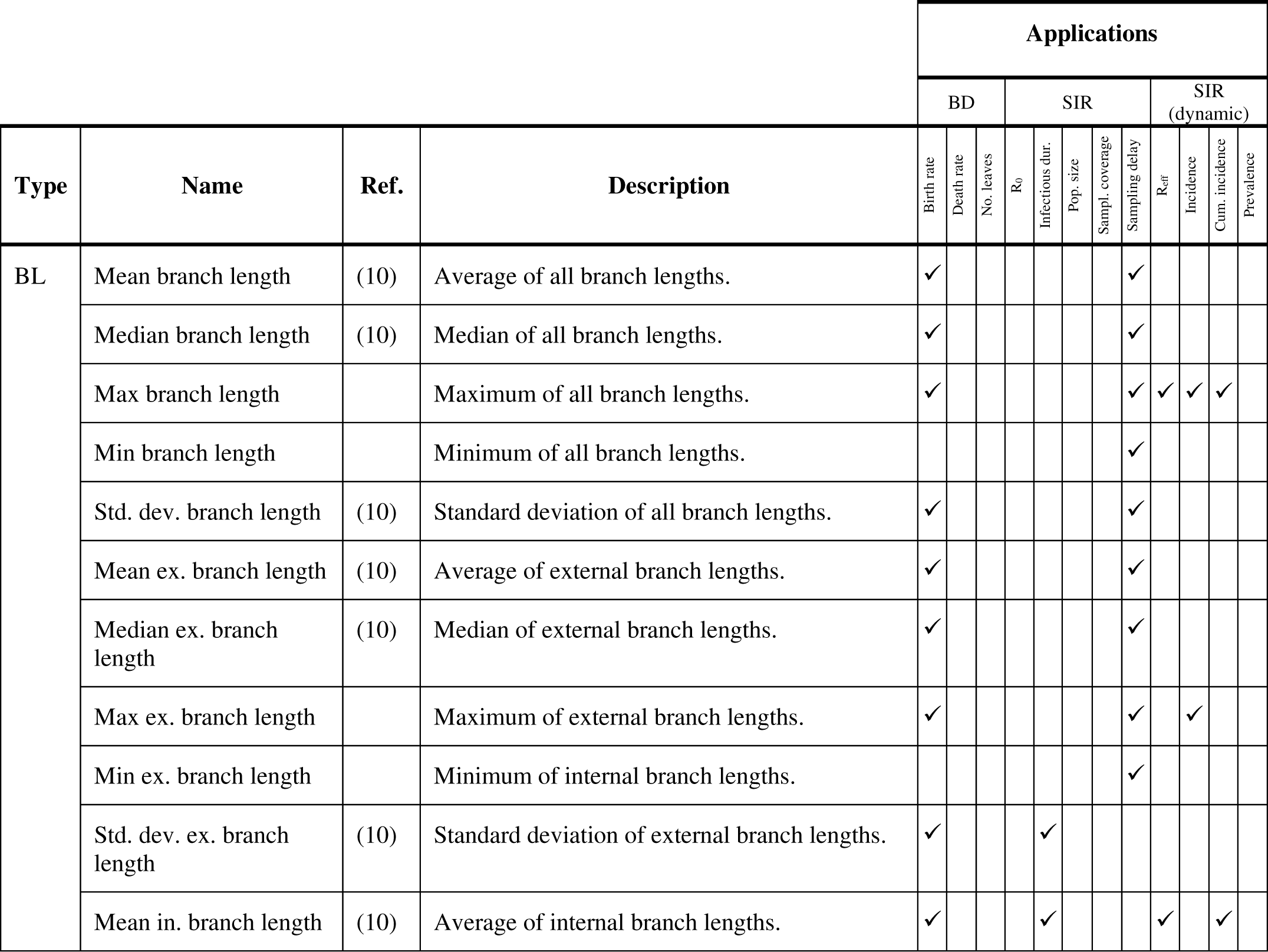
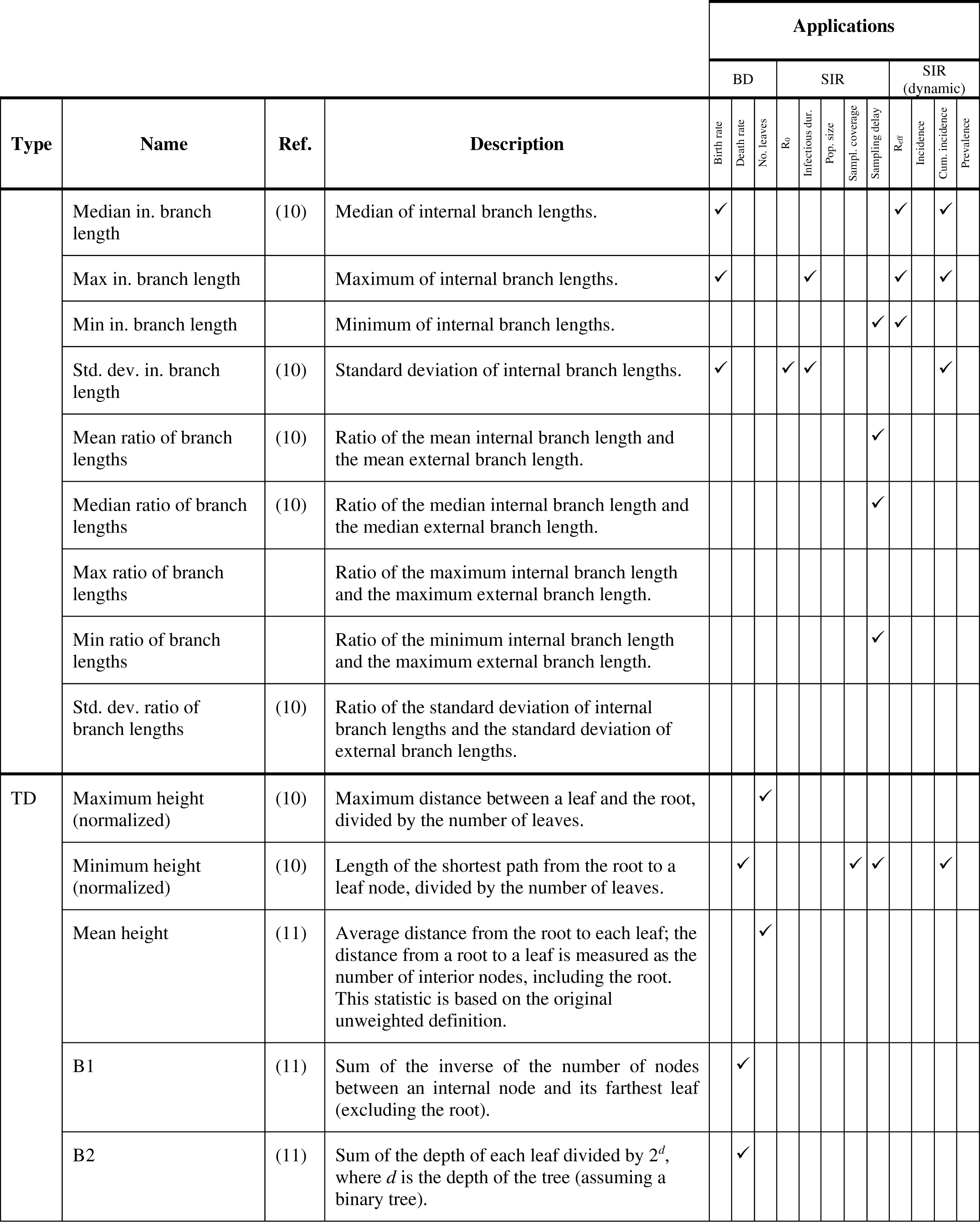
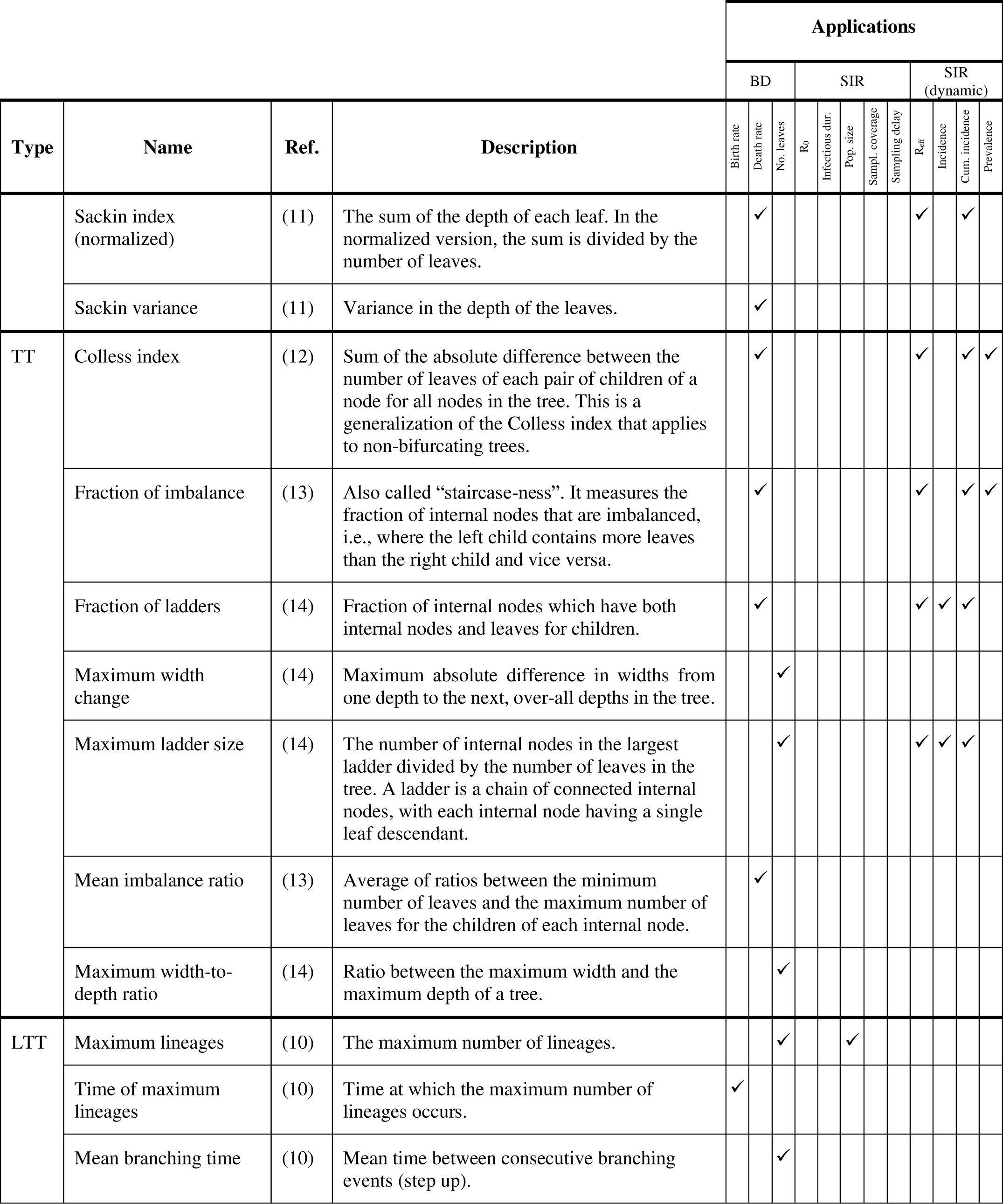
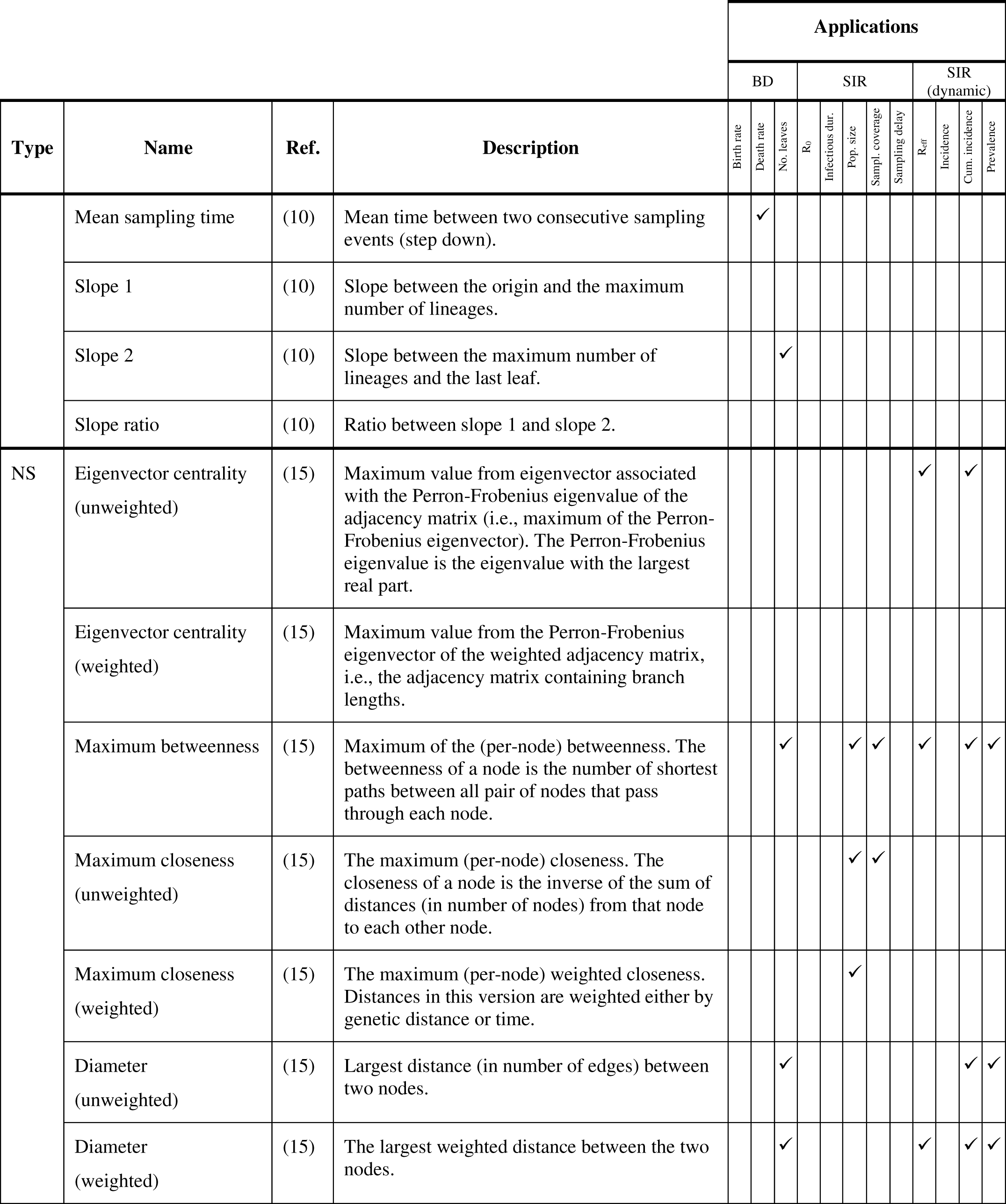
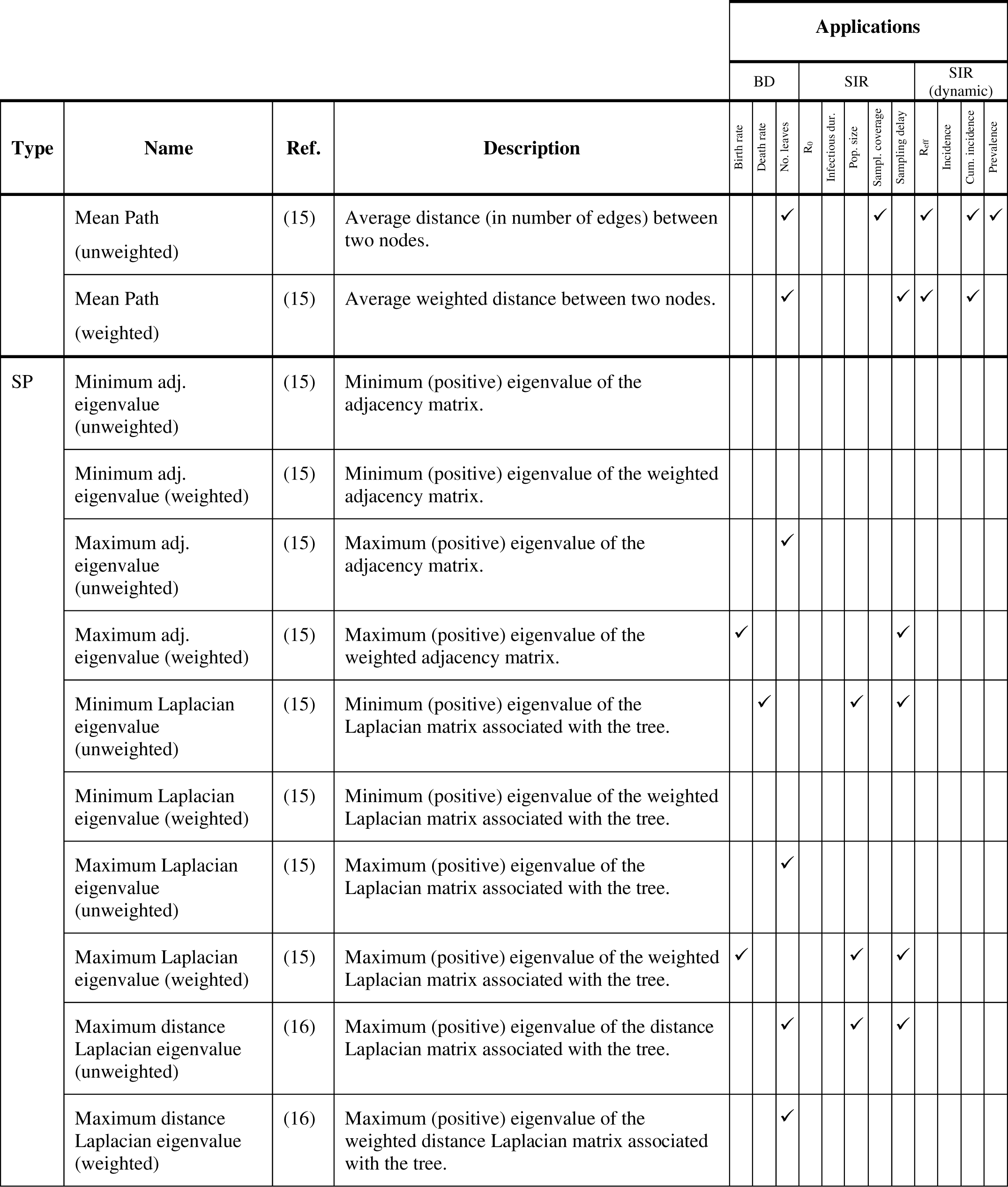
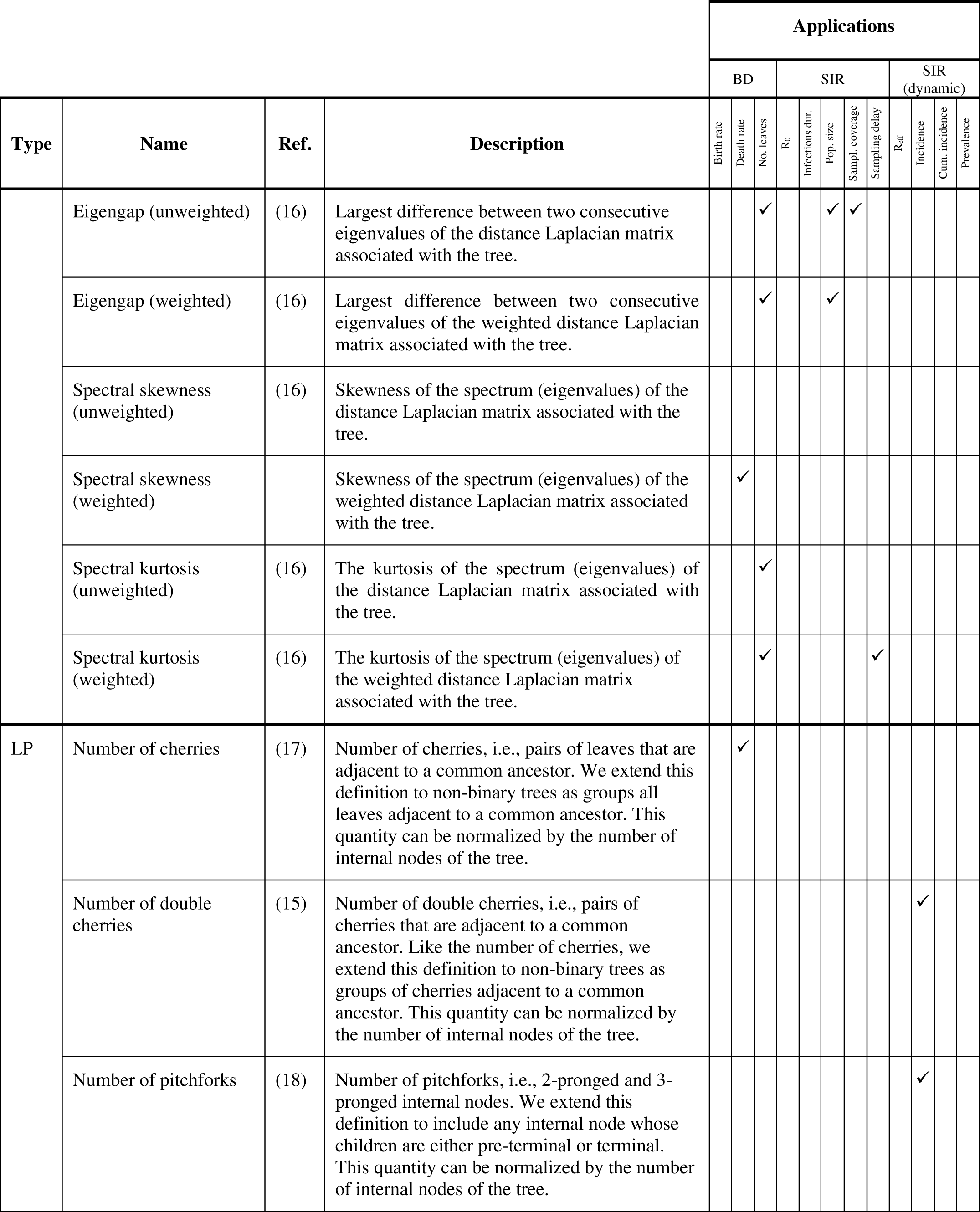
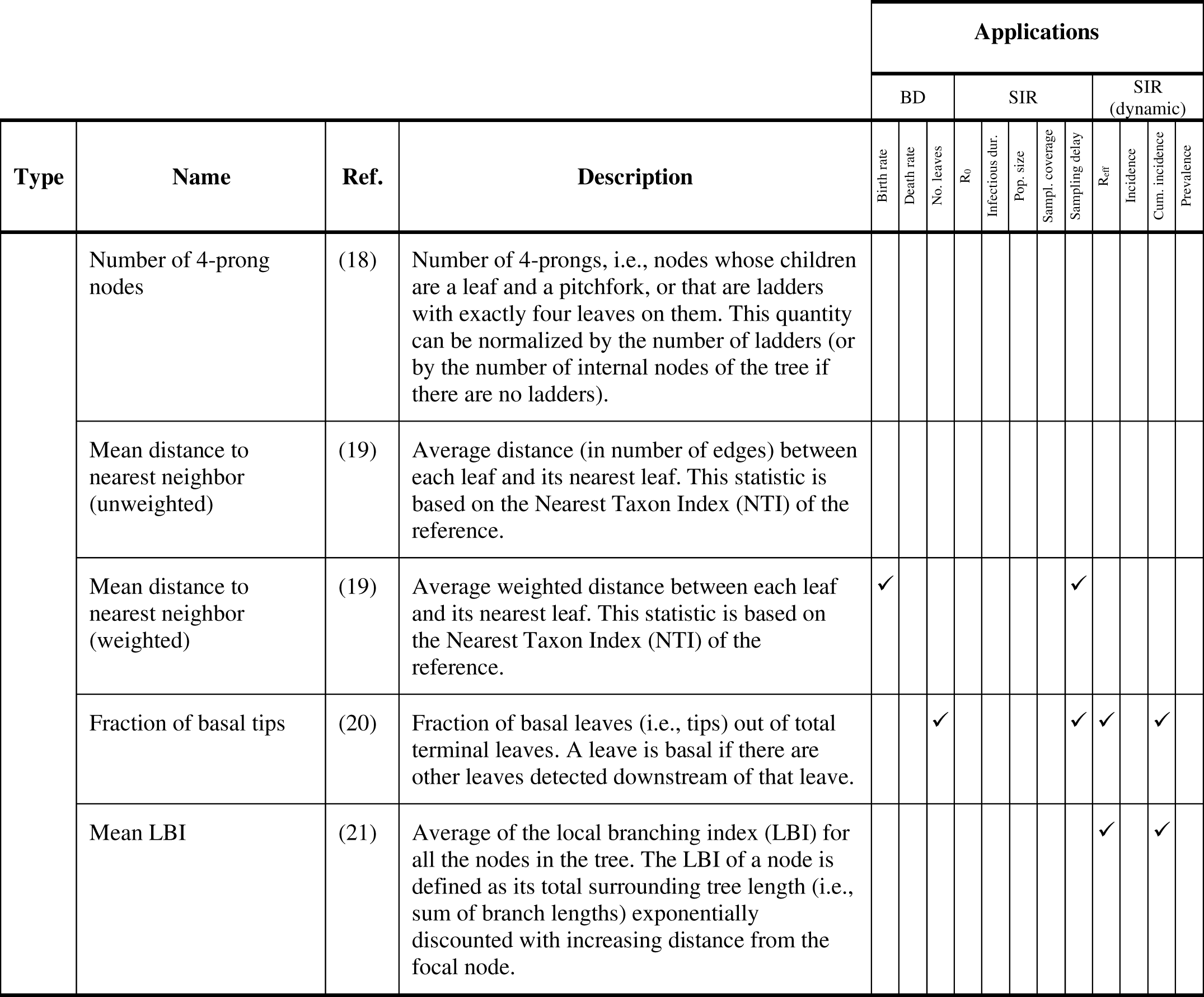
Summary statistics for phylogenetic trees. The types of summary statistics are: Branch Length (BL); Tree Depth (TD); Tree Topology (TT); Lineage Through Time (LTT); Network Science (NS); Spectral Properties (SP); and Local Properties (LP). The applications of each statistic are selected based on the correlation of the statistic with a parameter in one of our benchmark scenarios, namely, the Birth-Death model (BD) and the Susceptible-Infectious-Recovered model (SIR). A parameter exhibiting a correlation coefficient ≥0.6 with a given statistic is considered an application of that statistic (except for R_0_, where we consider correlation coefficients >0.55).

### 1.2 Contributions

This work addresses key gaps and challenges in the field of genomic epidemiology. Through a thorough literature review we identify a suite of phylogenetic summary statistics. Furthermore, we classify these summary statistics based on their properties, evaluate their performance in several benchmark scenarios, and quantify their computational performance. The benchmark scenarios evaluate the ability of summary statistics to identify changes in branching patterns through a birth-death model, and evaluate how well they estimate and track basic epidemiological quantities in a Susceptible-Infected-Recovered (SIR) disease transmission model. These benchmarks should be seen as an initial set of experiments, which could be complemented in future work by additional scenarios that evaluate the performance of the statistics based on the underlying network structure, sampling, local features of the trees, or their model calibration utility. Our work introduces an open-source python library, `phylomodels`, comprising functions and utilities tailored for phylogenetic workflows, which are designed to aid in the selection of informative metrics for epidemiological inquiries and model calibration.

## 2. Materials and Methods

### 2.1 Definitions

A phylogeny, or phylogenetic tree, is a graphic representation of the hypothesized evolutionary relationships among various biological strains or entities, based on similarities and differences in their genetic characteristics. A phylogeny shows which taxa are more closely related than others. Each leaf (or terminal node) represents an extant taxon, while an internal node signifies a common ancestor from which two or more descendants have evolved. Internal nodes may be called “hypothetical taxonomic units” to emphasize that they are hypothetical progenitors of leaves. Branches (or edges) connect nodes, representing evolutionary pathways. Branch lengths (patristic distances) denote the amount of evolutionary change or time between two nodes. Tree depth or tree height refers to the longest path from the root node to any leaf, indicating the maximum number of edges traversed. Tree width describes the maximum number of nodes at any level of the tree. For additional definitions of phylogenetic trees we refer the reader to (9).

From a graph theory perspective, a phylogenetic tree is a type of graph. Specifically, a phylogenetic tree is a connected acyclic graph. The adjacency matrix is a square matrix used to represent a finite graph, with rows and columns corresponding to the nodes (both internal and terminal) of the tree. The elements of this matrix indicate whether pairs of nodes are adjacent (connected by an edge), typically denoted as 1 for adjacency and 0 for non-adjacency. In phylogenetic trees, these entries can also represent the distance (either genetic or in time) between nodes. In this case, we may refer to it as a weighted adjacency matrix. The Laplacian matrix is derived from the adjacency matrix, highlighting the degree of each node and providing insights into the graph’s connectivity and structure. Specifically, the Laplacian matrix is defined as *L* = *D* – *A*, where *D* is the degree matrix (a diagonal matrix indicating the number of edges connected to each node) and A is the adjacency matrix. The Laplacian distance matrix has off-diagonal entries as the negative distances between nodes and diagonal entries as the sum of the distances in the corresponding row, making the rows sum to zero.

### 2.2 Summary statistics for phylogenetic trees

We evaluated the performance of 74 summary statistics in two different scenarios. The summary statistics, as well as their most useful applications (based on the benchmark experiments) are described in Table 1. The summary statistics have been implemented in `phylomodels`, which includes, in addition to efficient implementations of these summary statistics, other functions that are relevant for phylogenetic analysis such as tree reconstruction. The `phylomodels` library can be found at: http://github.com/InstituteforDiseaseModeling/phylomodels.

### 2.3 Birth-death model

Our first benchmark is based on trees simulated from a birth-death model via the `ngesh` package (20). The trees generated with this model do not have any direct relation to a specific disease or disease transmission model. However, these trees allow us to evaluate the effects different characteristics of a tree have on summary statistics without having to include the complexities of disease transmission, sampling, gene sequence simulation, and phylogenetic reconstruction.

In a birth-death model, each node has a probability λ of spawning a new node (i.e., birth), or a probability μ of dying. This model assumes that only one of two possible events (birth/death) can occur during any one event. Both of these events follow a Poisson process; the expected waiting time for an event follows an exponential distribution. The expected waiting time to the next birth is exponential with parameter λ, and the expected waiting time to the next death event is exponential with parameter μ. The expected waiting time to the next event (of any sort) is exponential with parameter λ + μ, and the probability that the event is speciation is λ/(μ + λ) or extinction is μ/(μ + λ).

Figure 1 shows examples of trees generated with a birth-rate model. In the figure, we keep the death rate constant while changing the birth rate. This leads to a visible pattern: the proportion of extant nodes increases as the difference λ – μ increases. We expect some summary statistics to quantitatively capture this pattern.

**Figure 1.**
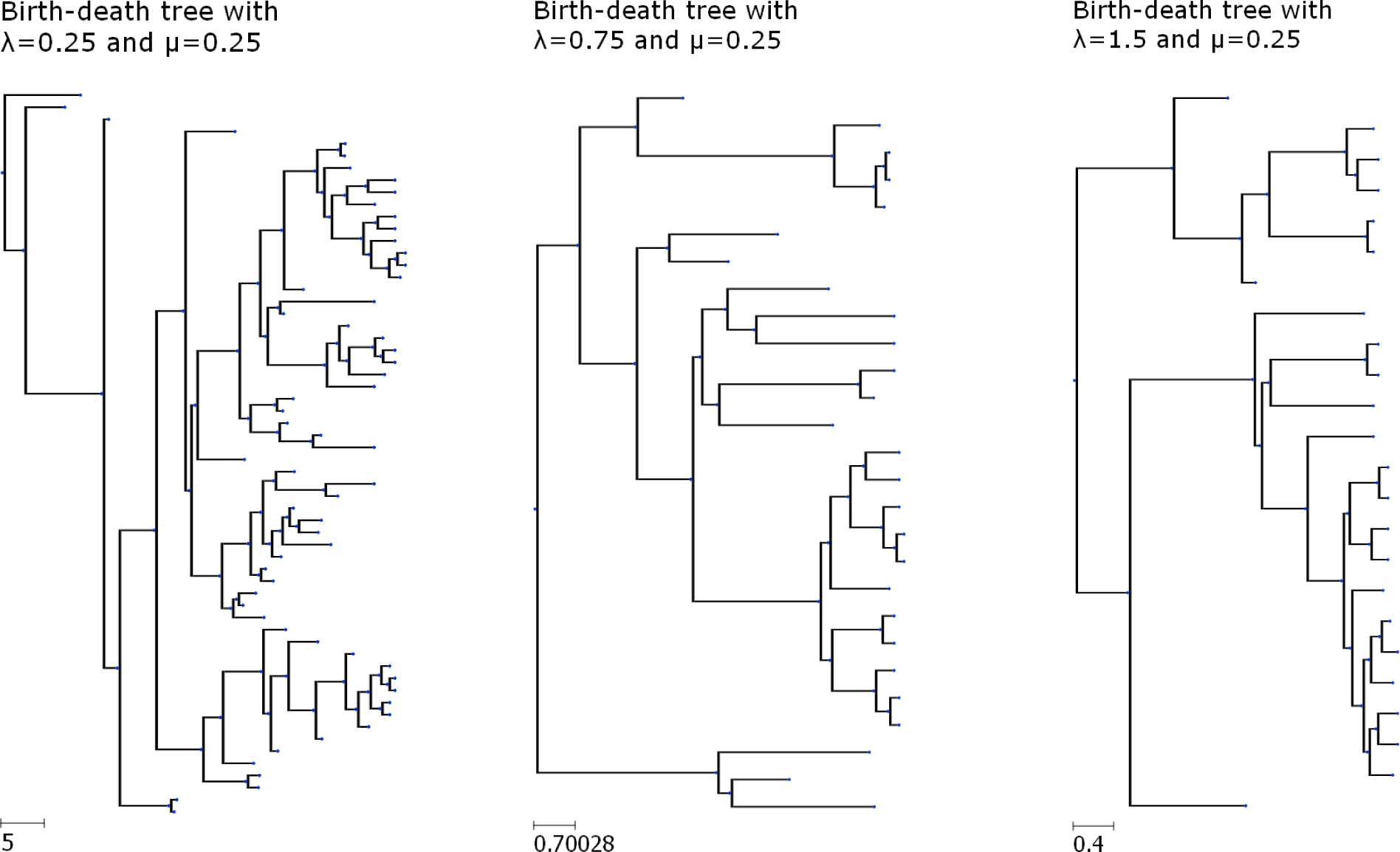
Examples of simulated phylogenetic trees generated using a birth-death model. In these trees, we change the birth rate while keeping the death rate constant. This leads to the trees exhibiting more extant nodes as the difference between birth and death rates increases. Note that the trees have different scales.

As a tool for characterizing the performance of a summary statistic, the birth-death model allows us to evaluate how well a summary statistic reacts to the rate at which nodes are being spawned (i.e., birth rate) as well as to the rate at which they stop spawning (i.e., death rate). In a disease transmission model, the birth rate could, for example, be a function of the reproductive number or transmission risk. Similarly, the death rate could be a function of factors such as the death or recovery rates.

For our analysis we used 40 evenly spaced values between 0.25 and 10 for both birth and death rate. For tree size we used 5 evenly spaced values between 50 and 250. Each point in parameter space was used five times for a total of 40,000 model runs.

### 2.4 Susceptible-Infected-Recovered (SIR) model

To characterize the summary statistics relationship not just to the tree but to the disease dynamics that created it, we utilized a simple agent-based model with SIR dynamics. In our SIR model, everyone begins as susceptible. Upon exposure to an infected individual, they transition to the infectious state and begin spreading the infection. The transmissibility of the disease is governed by the basic reproduction number (R_0_). An individual stays infectious for a specific period (infectious duration) before recovering and acquiring lifelong immunity (23).

Our software implementation of the SIR model uses a GPU to make computations significantly faster. The source code can be found at https://github.com/ghart-IDM/summaryStatisticsCharacterization/tree/main/SIR/gpu-SIR. In our model, the user can input a contact network, along with specific values of R_0_ and infectious duration. At each time step, every infected individual can infect each of their uninfected susceptible contacts. The probability of infecting another individual is a function of R_0_, the infection duration, and the number of contacts (i.e., the degree) of the (source) individual. The duration of infectiousness follows a Poisson distribution. The simulation ends when there are no more individuals in the infectious state. For our evaluations, we use a random network with constant degree. Each simulation starts with a single infection, and simulations with fewer than 11 infected individuals at the end are discarded.

The SIR model generates the transmission history represented as a line list. We then reconstruct the (simulated) phylogenetic tree. This process is depicted in Figure 2. It is based on three components, which are generated sequentially, as follows:

- **Line list.** The line list includes information about each infection event, namely, the source individual, the recipient (or newly infected) individual, the time of the transmission event, as well as the sampling time (i.e., the time when a sample would be collected from the newly infected individual). An example of a line list is shown in Figure 2(a).
- **Transmission tree.** The creation of a transmission tree based on this line list is straightforward. An example of a transmission tree for the line list in the figure is shown in Figure 2(b). In the transmission tree, nodes corresponding to infected individuals are placed at the time they were infected. The leaves in this tree represent individuals that did not transmit the infection, while internal nodes represent transmitters. Correspondingly, branch lengths in this tree represent the time between the transmitter acquiring the infection and a subsequent infection event. For example, in the figure, “B” was infected by “A” at time 4; subsequently, “B” infected “E” at time 10. Hence, “A”, which is a seed infection, sits at time 0 with a branch to “B” of length 4; similarly, “B” sits at time 4 with a branch to “E” of length 6.
- **Sampled phylogenetic tree.** Finally, we incorporate sampling times and convert the transmission tree into a phylogenetic tree. In the phylogenetic tree, all sampled individuals become leaves that are placed at the time their sample was taken. Internal nodes represent common ancestors placed at the time of their corresponding transmission events. This assumes that intra-host diversity is negligible. Internal nodes could also represent sampled non-transmitters appear at the time they are sampled. Their parent node represents the common ancestor at the time of infection, except for the case when the source individual is sampled before the transmission event. In that case, the parent node represents that (source) sampling event. Sampled transmitters that are sampled after their last transmission behave the same as non-transmitters, but their parent node is always at the time of their last transmission. Unsampled individuals are not in the phylogenetic tree. Note that, by simulating the tree in this way, we are assuming that the time reflected in the branch lengths is proportional to a mutation rate in scenarios where no recombination events occur.

**Figure 2.**
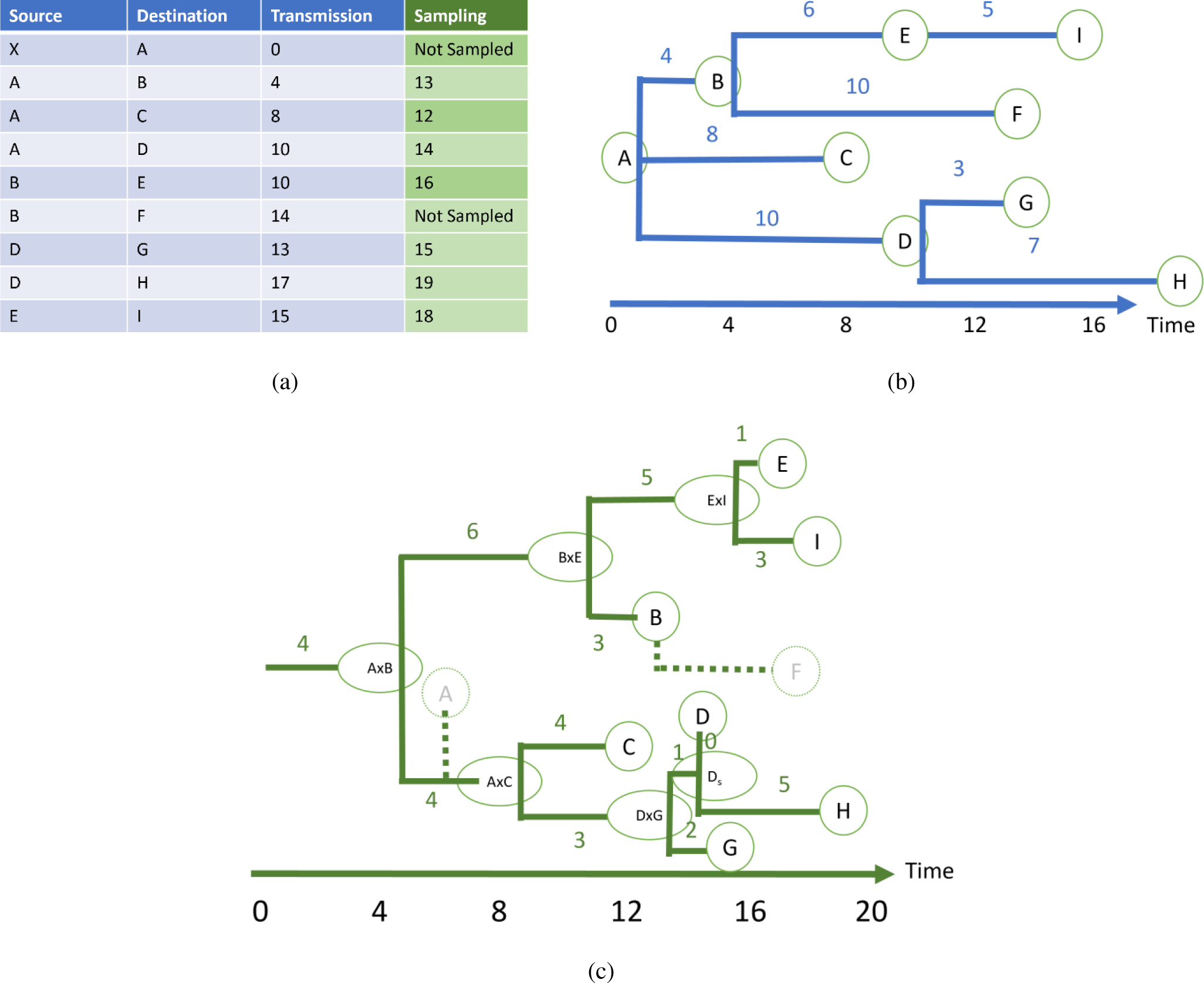
Illustration of the reconstruction process of a phylogenetic tree from transmission data. (a) Line list detailing source, destination, and transmission times for infections, as well as a sampling time for each infected individual. (b) Transmission tree visualizing time between transmission events. (c) phylogeny made from the transmission tree.

The reconstruction of the phylogenetic tree is implemented as a phylomodels utility that (1) builds the transmission tree, (2) converts it into a phylogenetic tree, and (3) samples from the phylogenetic tree.

For our analysis we used 6 evenly spaced values for the number of contacts ranging from 4 to 24, 6 evenly spaced values for the infectious duration ranging for 3 to 13 days, 6 evenly spaced values for R_0_ ranging from 2 to 12, 5 evenly spaced values for sample time ranging from 0 to 14 days, 5 evenly spaced value for sample fraction ranging from 0.2 to 1.0, and 5 evenly spaced values for population size ranging from 50 to 250. Each point in parameter space was evaluated 5 times, giving a total of 135,000 model runs.

### 2.5 Dynamic model

A crucial role of genomic epidemiology involves identifying changes in epidemics, such as increasing incidence, evolving infectiousness, or emerging variants. Here, we evaluated the effectiveness of summary statistics to track these dynamic values. Using our SIR model, we computed metrics such as the effective reproduction number (R_eff_), incidence, cumulative incidence, and prevalence on a weekly basis during an outbreak. We then calculated 40 time-dependent summary statistics, tailored to the specific characteristics of each week’s data, and assessed their correlation with dynamic model parameters across lag intervals from -2 to 2 weeks. The time lag with the highest correlation was identified and used, with most statistics exhibiting the strongest correlation at no lag. None of the statistics showed better correlations at time lags exceeding one week.

### 2.6 Evaluation of computational performance

Computational performance is a metric that should be considered when deciding whether to use a summary statistic or not. A summary statistic is not useful if its computation time is intractable for the size of the trees that need to be analyzed. Additionally, if two summary statistics can capture a tree’s property with similar accuracy, opting for the less computationally intensive statistic when computing it multiple times is preferable.

We assessed the computational performance of a summary statistic by measuring its execution time across tree sizes ranging from 25 to 6,400 leaves. The execution time was obtained by averaging 100 repetitions of each summary statistic for every tree size. All experiments were conducted on the same system –a system featuring an Intel Xeon Gold 6348 processor (16 cores at 2.60 GHz) with 64 GB of RAM. Although most summary statistics do not leverage parallelization, the matrix operations utilized by network science and spectral statistics fully use the available 16 cores. Theoretical complexity was also taken into account for each summary statistic where feasible.

## 3. Results

### 3.1 Correlation with epidemiological parameters

One direct application of summary statistics is to serve as proxies for parameters in a model or specific characteristics of disease transmission. We can assess the effectiveness of a summary statistic in this context by examining the correlation between the statistic and the model parameters. Here, we analyzed the correlation between the statistics and the parameters of our birth-death and SIR models.

The correlation between the summary statistics and the parameters in our birth-death model is illustrated in Figure 3. The columns in the figure represent the model parameters (i.e., birth rate, death rate), tree-size (indicated by the number of leaves), and the multiple correlation index. The rows correspond to the summary statistics. We can see that a significant portion of the branch-length summary statistics show a strong negative correlation with the birth rate, as changes in the birth rate led to variations in branch lengths, with higher birth rates resulting in shorter branch lengths (i.e., reduced time between spawning events). Some local and spectral statistics also exhibit high correlations with the birth rate. In total, 15 statistics show a strong correlation with the birth rate (considered strong correlation if the magnitude is greater than or equal to 0.7). Conversely, nine statistics correlate well with the death rate, with none of them falling under the branch-length category. None of the summary statistics demonstrate a strong correlation with both birth and death rates simultaneously. However, 32 statistics display a high multiple correlation value, indicating their effectiveness in capturing changes in both birth and death rates when provided together. Regarding tree size, 10 statistics exhibit a strong correlation with it, none of which belong to the branch-length type of summary statistic.

**Figure 3.**
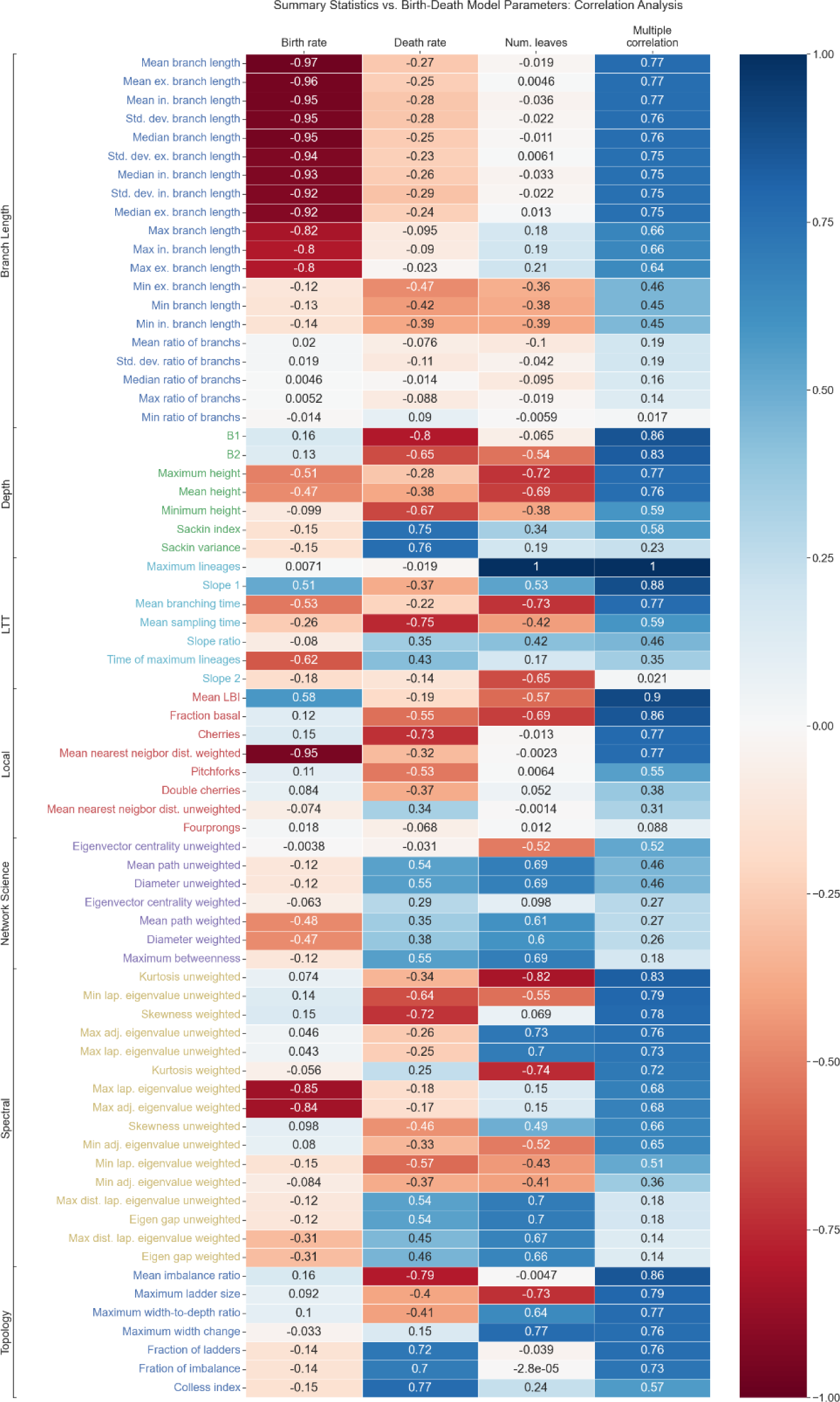
Correlation of statistics with parameters in the birth-death model. The statistics are grouped by their class/type, as described in Table 1. Blue-shaded squares indicate positive correlation, while red-shaded squares indicate negative correlation. The intensity of the color in each square reflects the magnitude of the correlation (the more intense the color the stronger the correlation). Several statistics demonstrate strong correlation with the birth-death model parameters and the resulting tree size, underscoring their utility for estimating these parameters in cases where their values are unknown.

Unlike the birth-death model, significant correlations are rarely observed in the SIR model. This is evident in Figure 4, which depicts the correlation between the statistics and parameters (both epidemic and sampling) of the SIR model, the tree size, and the coefficient of multiple correlation. Except for the coefficient of multiple correlation, correlations larger than 0.7 are infrequent. However, there are several instances of moderate to strong correlations (i.e., correlations with magnitudes greater than or equal to 0.4). Specifically, there are 22 statistics that show mild correlation with R_0_ (the basic reproductive number), 20 with the duration of infection, 26 with the total population size, 25 with sample coverage, and 24 with sampling delay. Among all the correlations with R_0_, number of contacts, infectious duration, population size, and sampling coverage, the highest value was observed for the correlation between the standard deviation of external branch lengths and the infectious duration. None of the statistics show correlation with the number of contacts, suggesting that these summary statistics are not ideal for directly estimating the number of contacts in an underlying transmission network. Conversely, in the case of sampling delay, a considerable number of statistics (a total of 16) demonstrate correlation magnitudes exceeding 0.7. This indicates that many of these statistics respond to changes in sampling delay, which could significantly impact the structure of trees.

**Figure 4.**
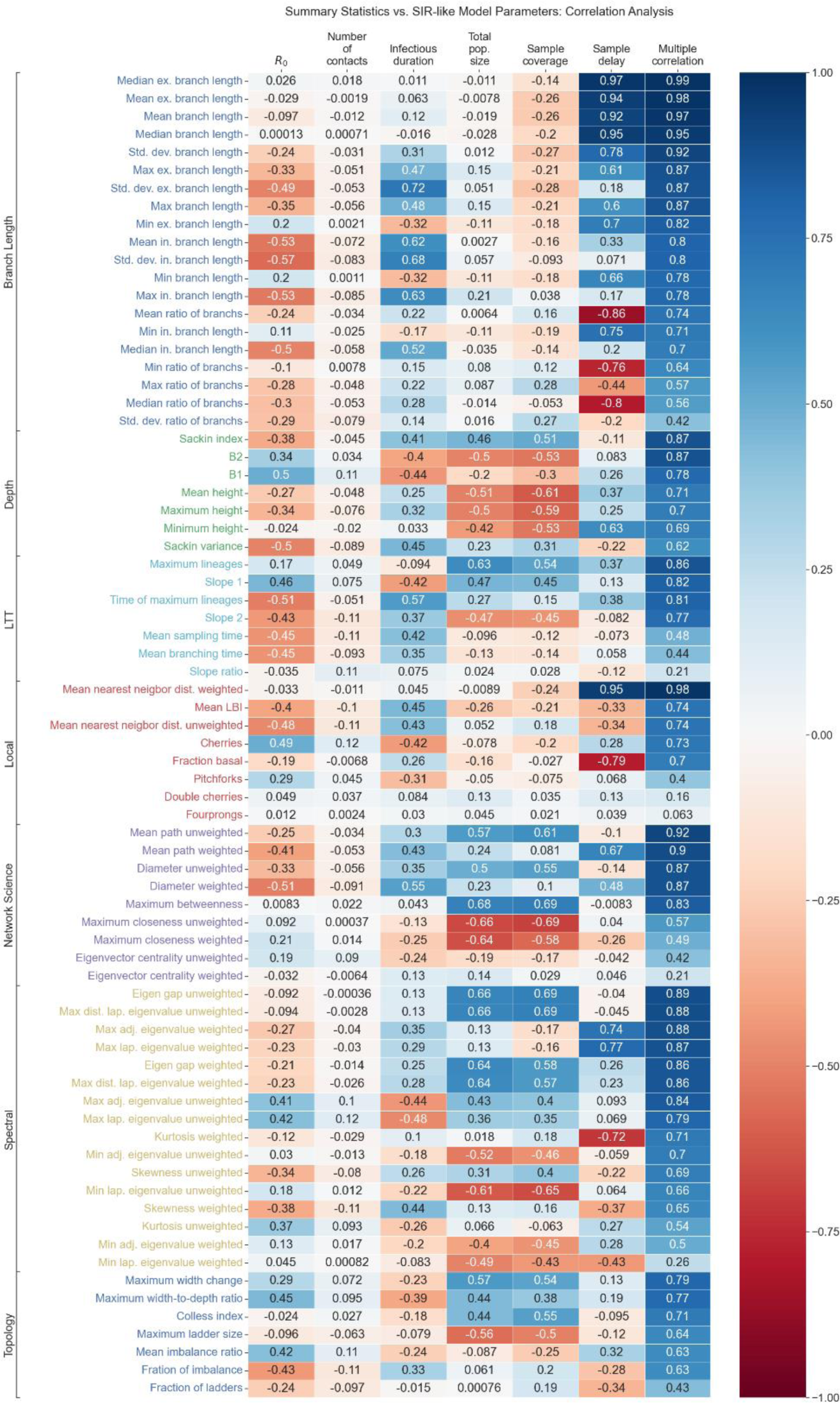
Correlation of statistics with parameters in the SIR model. The statistics are grouped by their class/type, as described in Table 1. Blue-shaded squares indicate positive correlation, while red-shaded squares indicate negative correlation. The intensity of the color in each square reflects the magnitude of the correlation (the more intense the color, the stronger the correlation). Several statistics demonstrate significant correlation with the SIR model and sampling parameters, underscoring their utility for estimating these parameters in cases where their values are unknown.

An interesting scenario arises when examining the dynamic characteristics of the SIR model. As mentioned in Section 2.5 above, certain parameters or outcomes undergo changes as an epidemic unfolds. For instance, the effective reproductive number (R_eff_), incidence, cumulative incidence, and prevalence, evolve as an epidemic advances within a specific population. We analyzed the correlation of these parameters with our set of summary statistics to gauge how effectively these statistics respond and adjust to variations in the epidemic model parameters. Specifically, we assessed the correlation between the time series of each of the epidemic parameters or outcomes, and the time series of each statistic. The results are shown in Figure 5. We observe that 9 of the statistics exhibit strong (i.e., ≥0.7) correlation with R_eff_, and 15 of them show mild to high (i.e., between 0.4 and 0.7) correlation with the cumulative incidence. Interestingly, only the maximum ladder statistic displays strong correlation (equal to 0.76) with incidence.

**Figure 5.**
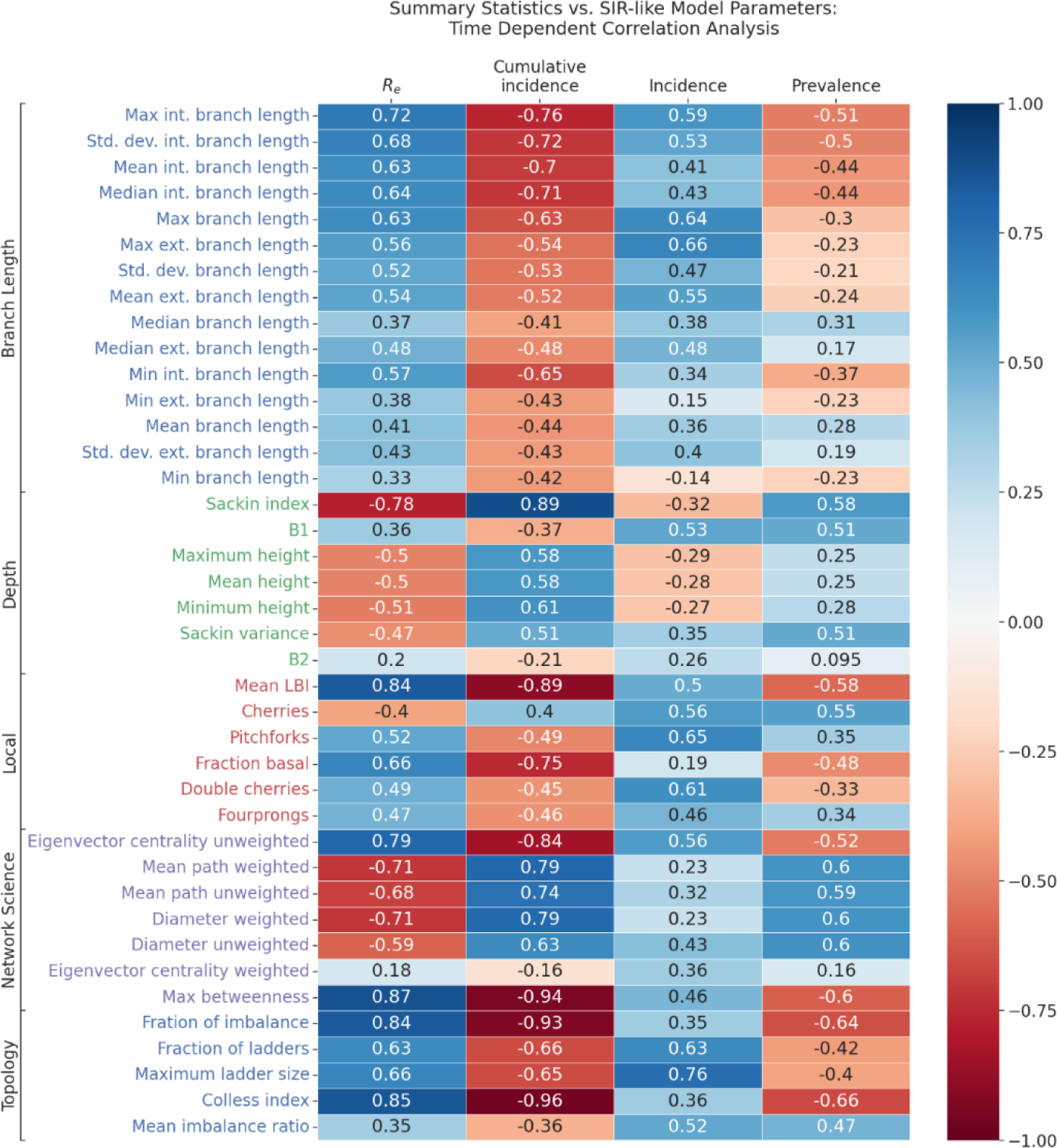
Correlation of statistics with dynamic parameters or outcomes of the SIR model. The statistics are grouped by their class/type, as described in Table 1. Blue-shaded squares indicate positive correlation, while red-shaded squares indicate negative correlation. The intensity of the color in each square reflects the magnitude of the correlation (the more intense the color, the stronger the correlation). Several statistics demonstrate significant correlation with the model parameters or outcomes, underscoring their utility for estimating these dynamic parameters in cases where their values are unknown.

Based on the correlation analysis conducted for our benchmark scenarios, we categorize each statistic as useful or effective for estimating a particular parameter if its correlation with that parameter is equal to or greater than 0.6. To facilitate assessment, we present these findings in the final column of Table 1, which provides a comprehensive list and description of each statistic. Overall, it is evident that multiple statistics can be utilized for estimating a specific parameter. This underscores the necessity for guidelines to aid in determining the appropriate statistic(s) for a given application. Further insights on this matter are elaborated in Section 4 below.

### 3.2 Computational performance

As mentioned above, computational performance is an important factor when considering the use of a summary statistic. This is particularly crucial when repetitive computation of a summary statistic is required, as is seen in scenarios like model calibration. It also serves as a criterion for favoring less computationally intensive summary statistics when multiple options exhibit similar accuracy or proficiency in quantifying phylogenetic tree’s characteristics.

When looking at the summary statistics analyzed in this study, it is evident that a significant portion of them exhibit linear scalability relative to tree size. These statistics primarily involve reading each node or branch of the tree just once. This category encompasses statistics related to branch length, depth, local, lineage-through-time, and topology. In contrast, statistics from network science or spectral groups demonstrate exponential scalability ranging from *O*(N^2^) (i.e., quadratic scalability) to *O*(N^3^) (i.e., cubic scalability), with N being the number of nodes of the tree. This scaling behavior is shown in Figure 6, which compares the performance of various groups of statistics across trees of varying sizes. The figure exclusively considers summary statistics that exhibit strong correlation (correlation coefficients larger than or equal to 0.7) with the epidemiological parameters outlined in our benchmark scenarios. It is worth noting that branch length statistics show a roughly 3-4 times slower performance compared to other linearly-scaling statistics. This discrepancy could be attributed to the increased data access requirements, specifically involving the branch weights.

**Figure 6.**
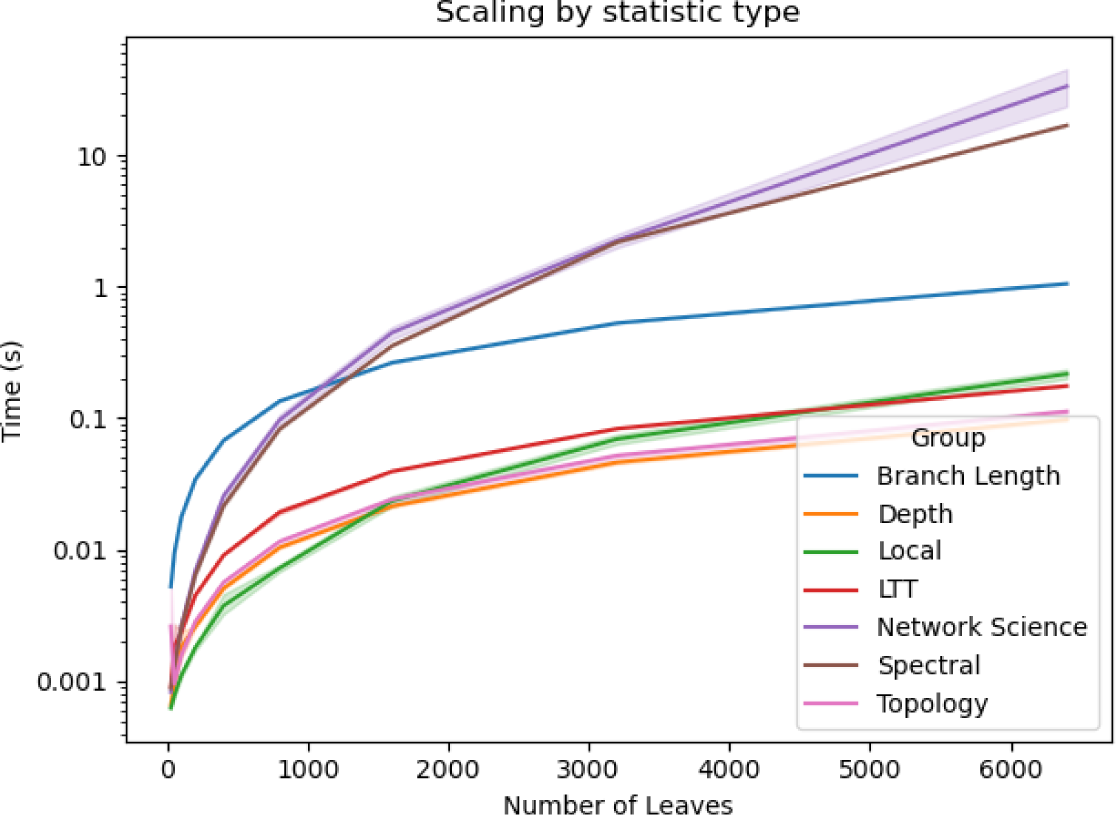
Time performance comparison across tree sizes for strongly correlated summary statistics. The results are grouped by the summary statistics’ categories.

Further details on the time performance of individual summary statistics are presented in Figure 7. Notably, summary statistics categorized within the local, network science, and topology groups, exhibit significant variability within each group. Conversely, summary statistics from other groups demonstrate consistent behavior within their respective groups, indicating that the computational procedures involved in computing these statistics are quite similar. Across all cases, it is evident that the time required to compute a summary statistic increases proportionally with the tree’s size, emphasizing that none of the summary statistics maintain a constant computation time.

**Figure 7.**
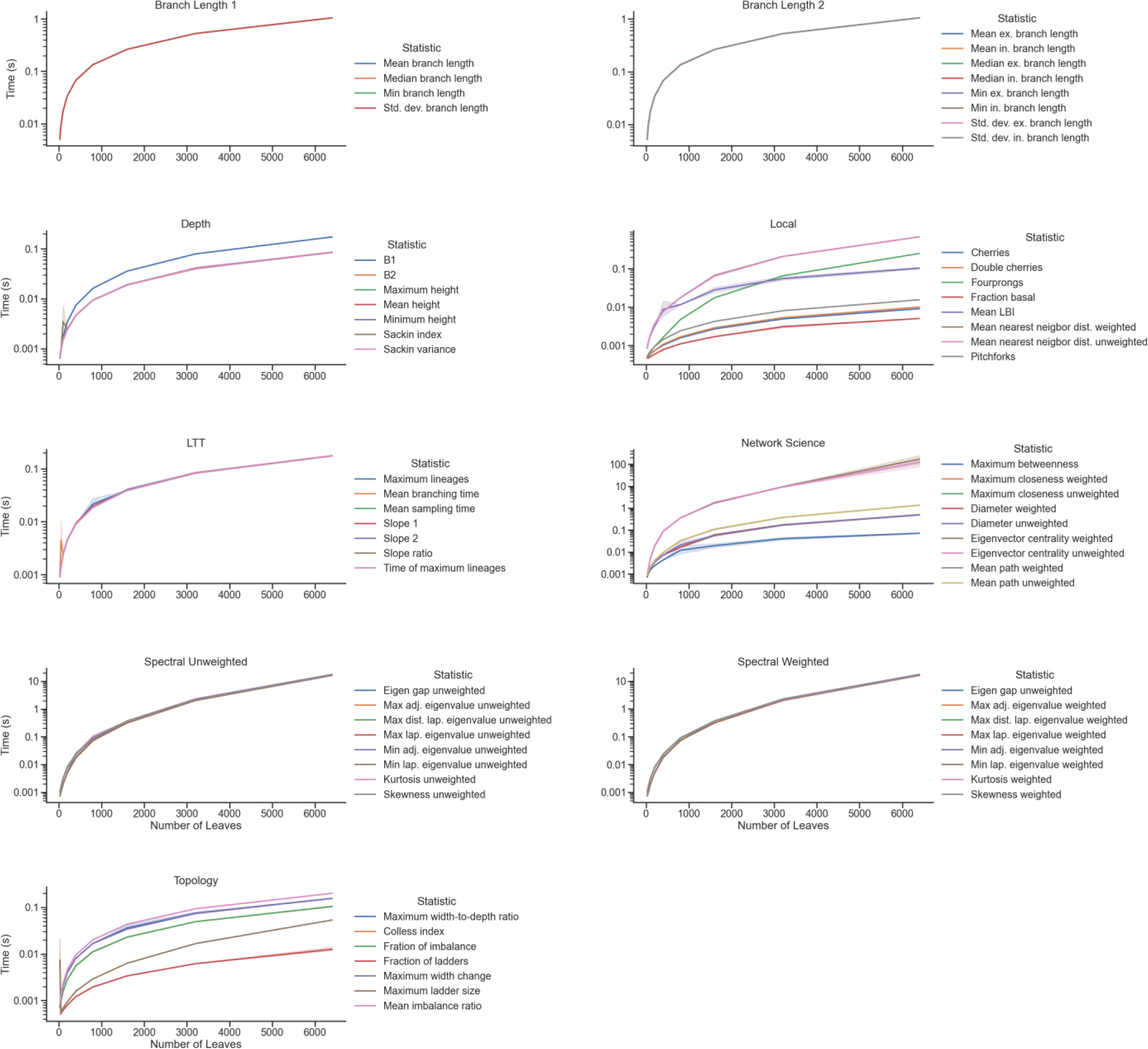
Time performance comparison of summary statistics across tree sizes. The figures display results for groups of statistics within distinct categories: branch length, depth, local, lineage-through-time, topology, network science, and spectral.

Among the summary statistics, the fastest to compute include cherries, double cherries, pitchforks, fraction basal, and fraction of ladders (see definitions in Table 1 above). Note that most of these fast-performing statistics belong to the local tree properties group. On the other end of the spectrum, the most computationally demanding summary statistics are the weighted and unweighted eigenvalue centralities (see definitions in Table 1 above), which fall under the network science category. Notably, eigenvalue centrality’s computation time is approximately one order of magnitude slower than all spectral summary statistics, which, in turn, are at least an order of magnitude slower than all other summary statistics evaluated in this study. This increased complexity arises from the need to compute a full eigenvalue decomposition with high precision. In contrast, other spectral summary statistics often involve more straightforward eigenvalue computations or approximations that are computationally less demanding than those involved in the computation of the eigenvalue centrality.

## 4. Discussion

The comprehensive analysis presented above enables us to discern the specific strengths and applications of each of the summary statistics that were subjected to evaluation. This analysis sheds light on their utility within the context of the benchmark scenarios outlined in this paper. The results of our evaluation, delineating the unique strengths and purposes of each summary statistic, are presented in the final column of Table 1. While most summary statistics have demonstrated utility in various contexts, the diverse array of statistics, parameters, models, and problems of interest necessitates a nuanced understanding of their respective applications. Building upon these findings, we have identified several additional critical observations, as follows:

▪ **The choice of an optimal tree summary statistic depends on the specific application or property under consideration for estimation.** While this may seem self-evident, it is a crucial factor that warrants careful consideration. Through our analysis, we have identified statistics that excel in estimating parameters within our benchmark scenarios. For instance, the local branching index was particularly useful for assessing changes in the effective reproductive number. Similarly, several branch-length summary statistics demonstrated the ability to quickly estimate sampling delays. While these findings can directly inform applications closely aligned with these scenarios, it is essential to recognize that varying contexts may necessitate additional scrutiny and potentially the development of novel statistics to enhance accuracy and efficacy in estimation outcomes. Some scenarios may even call for the design of new summary statistics. For example, if one needs to estimate the number of contacts based on tree data, none of the summary statistics described here will work, and a new one should be created. These observations underscore the importance of tailoring the choice of statistics to the unique requirements of each application, ensuring optimal performance and robust results.
▪ **The performance of statistics in estimating specific parameters does not necessarily extend to their ability to estimate other variables or parameters.** In other words, not all statistics that excel in estimating one parameter or quantity are equally proficient in estimating other variables or parameters. This observation underscores the nuanced nature of statistical estimation within the context of phylogenetic trees. For instance, eigenvector centrality correlates well with effective reproductive number and cumulative incidence, but not with other metrics such as incidence, sampling delay, or infection duration. Similarly, the number of double cherries and pitchforks correlated well with incidence, but not with effective reproductive number, prevalence, cumulative incidence, or sampling delay. This highlights the necessity of employing multiple statistics when aiming to estimate a diverse range of variables accurately.
▪ **Choosing between multiple statistics of similar effectiveness can be a critical task.** When presented with several statistics that show equivalent efficacy in estimating a particular quantity, parameter, or property, the selection process becomes extremely important. While opting for a statistic with potentially higher correlation may seem intuitive, it is essential to avoid redundancy by employing multiple highly correlated statistics to estimate the same parameter. Such redundancy not only consumes unnecessary resources but also fails to enhance estimation accuracy significantly. In such scenarios, factors such as computational complexity, execution time, accuracy, and reliability measures should be carefully evaluated to guide the selection of the most appropriate statistic. Considering these aspects can aid in determining the optimal statistic to utilize when confronted with multiple viable options. Additionally, the adaptability of statistics to changes in the scenario should also be taken into account, as this can influence the robustness and efficiency of the estimation process.
▪ **We need to balance the information delivered by spectral and network science statistics with their computational complexity.** The utilization of summary statistics originating from network science or relying on spectral measures can potentially offer a wealth of information that may be crucial for certain analyses. However, it is prudent to exercise caution and reserve their application for cases where the desired information cannot be adequately captured by simpler summary statistics. This cautious approach arises from the computational intensity associated with network science and spectral statistics, as they exhibit exponential scaling in relation to the size of the phylogenetic tree. When faced with the necessity of computing these statistics repeatedly, users must prioritize employing optimized implementations to mitigate the computational burden effectively. Alternatively, investing efforts in optimizing the computation process can prove instrumental in enhancing efficiency and ensuring the practicality of employing these complex statistics in phylogenetics analyses.
▪ **For model calibration and repetitive tasks, use the fastest summary statistics among the highly correlated ones.** This rule of thumb is supported by our experimental findings, where we observed that multiple summary statistics can effectively capture a property of interest, with most scaling linearly in tree size. However, certain statistics exhibit significantly longer computation times (1 to 2 orders of magnitude) and additional overhead than others. For example, when prevalence is the calibration target, several summary statistics correlate well with it, such as Colless index, fraction of imbalance, maximum betweenness, and weighted/unweighted mean path. However, computing the mean path can take an order of magnitude more time than computing any of the other quantities, so it should be discarded due to its complexity. Furthermore, using multiple correlated statistics is unnecessary for accuracy, as this practice does not enhance overall precision but rather increases computational complexity in phylogenetic analyses. In the previous example, our findings showed that the Colless index had the strongest correlation with prevalence, making it the preferred summary statistic for use in the example calibration job scenario.

## 5. Conclusions

The field of genomic epidemiology stands at the forefront of modern public health research, promising to revolutionize our approach to disease surveillance and control. However, it faces significant challenges related to the computationally intensive nature of phylogenetic analysis, the interpretation of complex genetic data, and the integration of these findings with traditional epidemiological information. Our work aims to address these challenges by providing a robust framework for the strategic selection of summary statistics that are specifically tailored to meet the needs of epidemiological investigations and model calibration. This framework begins with two benchmark scenarios, namely, a birth-death process, and an SIR model. By conducting a systematic evaluation of multiple summary statistics in simulated outbreak scenarios, we offer valuable insights into the selection of informative metrics that strike a delicate balance between computational efficiency and analytical accuracy. Through this comprehensive study, we not only deepen our understanding of the intricate dynamics of disease spread but also establish practical guidelines for researchers and practitioners to leverage phylogenetic data effectively. To further support this endeavor, we have developed a Python library called `phylomodels`, which provides the summary statistics analyzed in this paper, as well as additional utilities for phylogenetic workflows, thereby enhancing the tools available to the genomic epidemiology community.

## Acknowledgements

This work is based on research conducted by Institute for Disease Modeling, a research group within, and solely funded by, the Bill & Melinda Gates Foundation.

We would like to express our sincere gratitude to Mandy Izzo for her invaluable comments, revisions, and editorial assistance. Her meticulous work and insightful feedback have significantly enhanced the quality of this manuscript.

## Code Availability

All code used for this analysis can be found at https://github.com/ghart-IDM/summaryStatisticsCharacterization/tree/main and the libraries mentioned in the text ngesh and phylomodels

## Notes

https://github.com/ghart-IDM/summaryStatisticsCharacterization

### Competing Interest Statement

The authors have declared no competing interest.

